# The molecular structure of an axle-less F_1_-ATPase

**DOI:** 10.1101/2024.08.08.607276

**Authors:** Emily J. Furlong, Ian-Blaine P. Reininger-Chatzigiannakis, Yi C. Zeng, Simon H. J. Brown, Meghna Sobti, Alastair G. Stewart

## Abstract

F_1_F_o_ ATP synthase is a molecular rotary motor that can generate ATP using a transmembrane proton motive force. Isolated F_1_-ATPase catalytic cores can hydrolyse ATP, passing through a series of conformational states involving rotation of the central γ rotor subunit and the opening and closing of the catalytic β subunits. Cooperativity in F_1_-ATPase has long thought to be conferred through the γ subunit, with three key interaction sites between the γ and β subunits being identified. Single molecule studies have demonstrated that the F_1_ complexes lacking the γ axle still “rotate” and hydrolyse ATP, but with less efficiency. We solved the cryogenic electron microscopy structure of an axle-less *Bacillus* sp. PS3 F_1_-ATPase. The unexpected binding-dwell conformation of the structure in combination with the observed lack of interactions between the axle-less γ and the open β subunit suggests that the complete γ subunit is important for coordinating efficient ATP binding of F_1_-ATPase.

## 1. Introduction

F_1_F_o_ ATP synthase is a molecular rotary motor that can generate ATP using a transmembrane proton motive force [1–3]. The F_1_ region (often termed the F_1_-ATPase) is a rotary ATPase composed of three catalytically active β subunits separated by three catalytically inactive α subunits, arranged in a hexameric ring around the N- and C-terminal helices of the central γ subunit (Fig. S1) [4]. In isolation the F_1_-ATPase can hydrolyze ATP inducing rotation of the γ subunit (generally referred to as the “rotor”) [5–7]. In the intact F_1_F_o_ complex the F_1_-ATPase is coupled to the F_o_ region that drives the translocation of protons across the membrane, allowing the synthesis of ATP by the F_1_-ATPase [6].

*Geobacillus stearothermophilus* (also known as *Bacillus sp.* PS3) F_1_ ATPase (termed TF_1_ for Thermophilic F_1_ ATPase) has been used as a model for understanding F_1_-ATPase hydrolysis activity [5, 8–10]. Experiments performed using single particle cryogenic electron microscopy (cryo-EM) [11] and single-molecule rotation studies [5, 12, 13] have facilitated a detailed mechanistic study of this enzyme undergoing ATP hydrolysis. One full catalytic cycle of F_1_-ATPase results in the hydrolysis of three ATP molecules, each of which is sequentially bound, hydrolyzed to ADP and P_i_, and released by the each of the three β active sites. Throughout catalysis, the β subunits cycle through different conformations (opening and closing) that generate rotation of the γ subunit rotor [4, 11]. Each ATP hydrolysis step drives a 120° the rotation of the γ subunit [4, 6, 10], so that after a complete catalytic cycle it has rotated a full 360°. Single molecule rotation studies have found that these 120° steps can be further divided into two sub-steps of ∼80° and ∼40° [13]. Cryo-EM structures of the TF_1_ in a “binding-dwell” conformation (prior to the ∼80° ATP binding stroke) and a “catalytic-dwell” conformation (before the ∼40° hydrolysis stroke) have shown the molecular details of how the β subunits open and close through a series of states during catalysis to induce rotation of the γ subunit [11]. These two conformations were captured using a temperature sensitive TF_1_ mutant (βE190D) that at 10°C spends significantly more time in the binding-dwell and at 28°C spends significantly more time in the catalytic-dwell, when incubated with MgATP [14].

For the F_1_-ATPase to operate as a multi-site rotary motor, the subunits must act cooperatively. Previous studies have suggested that the γ subunit is responsible for mediating cooperativity between the β active sites [15]. Three key interaction sites between the β and γ subunits have been identified: (1) the hydrophobic sleeve; (2) the β-catch loop; and (3) the β-lever domains (Fig. S2) [4, 16, 17]. In TF_1_ numbering, the hydrophobic sleeve includes residues α280-285 and β271-275 that surround residues 271-283 at the tip of the γ subunit’s C-terminal helix [4]. The TF_1_ β-catch loop refers to residues 312-315 of β that interact with residues 262-263 of γ [4]. The β-lever domains, residues β382-396, undergo the largest relative movement during catalysis and interact with the rotor at positions γ15-31 and γ243-253 [17].

Other studies have shown that the γ subunit is not essential for F_1_-ATPase activity, because rotorless TF_1_ (i.e. lacking a γ subunit) is still able to hydrolyze ATP, with the α/β subunits undergoing similar sequential conformational changes as the rotor containing F_1_-ATPase [18, 19]. However, both the rotation speed and hydrolysis activity of rotorless TF_1_ are reduced significantly [18] (∼60 times slower hydrolysis rate; ∼300 s^-1^ vs ∼5 s^-1^ [20]), showing that although it is not essential, the γ subunit rotor is required for efficient catalysis. Previous studies have investigated the role of the axle region of the γ subunit in TF_1_ rotation by truncating either or both of the N- and C-termini of γ and assessing the rotation rate using single molecule rotation analysis [21, 22]. Both rotation rate and stability of the complex decreased as the truncation increased, with the mutant lacking the entire axle (γΔN22C43) behaving similarly to the rotorless mutant [20, 21].

To investigate the role of the γ axle in efficient catalysis in greater detail, we have solved the structure of the TF_1_ γΔN4C25 mutant [21] (referred to here as axle-less TF_1_) using single particle cryo-EM. The structure reveals that although this mutant cannot interact with the hydrophobic sleeve or the β-catch loop, enough of the axle remains to form contacts with the β-lever domains. Additionally, the structure of the TF_1_ γΔN4C25 mutant exhibits an unexpected binding-dwell conformation that provides insight into why the full-length axle is required for efficient catalysis.

## 2. Results and discussion

### 2.1 Purification and activity of axle-less TF_1_

The γ subunit of the axle-less TF_1_ examined in this study was truncated by four amino acids from the N-terminus and twenty-five amino acids from the C-terminus (γΔN4C25, Fig. 1A). This mutant was chosen as it was predicted to result in the loss of both the hydrophobic sleeve and β-catch loop interactions whilst maintaining enough of the γ axle to form a stable complex with the α and β subunits. Previous work [21] and preliminary cryo-EM studies showed that standard His-tag purification strategies of γΔN4C25 TF_1_ resulted in a significant amount of rotorless TF_1_ contamination, highlighting the potential instability of axle-less TF_1_. As rotorless TF_1_ would likely interfere with cryo-EM data processing and bulk activity measurements, we adopted a dual affinity purification strategy and, in addition to the N-terminal His-tag on the β subunit, a strep-tag II was inserted into a loop region of the γ subunit (between residues N196 and K197) (Fig. 1A). Dual affinity purification together with a final size exclusion chromatography step resulted in purified axle-less TF_1_ that remained mostly intact and stable after one freeze-thaw cycle (Fig. 1B, S3A). ATP regeneration assays were used to assess the hydrolysis activity of the axle-less TF_1_, compared to the wild type (WT) TF_1_ (note the WT used in this study had mutations αC193S+W463F and γS109C+I212C) and showed that the axle-less mutant was approximately 10-fold less active than the WT, consistent with previous findings [21] (Fig. 1C).

**Fig. 1.**
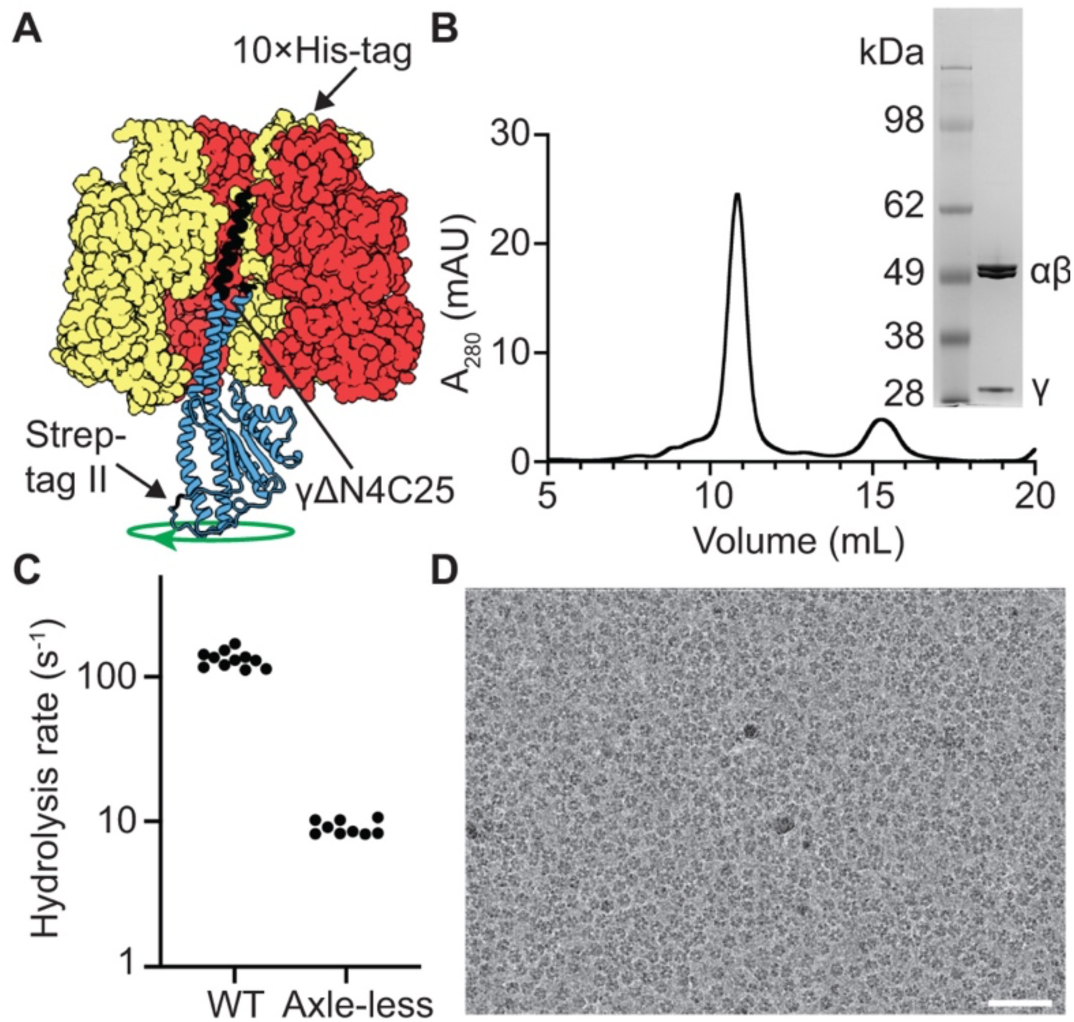
Purification and ATPase activity of axle-less TF_1_. (**A**) Structure of TF_1_ showing the truncation made to the γ axle (black) and the locations of the affinity tags used for purification. The remaining portion of the γ subunit is shown in blue, β subunits are yellow, and α subunits are red. One α and one β subunit have been removed to make the γ subunit visible. Green arrow depicts the direction of rotation during hydrolysis (counterclockwise when viewed from below/the membrane). (**B**) Size exclusion chromatogram and SDS-PAGE of the purified axle-less TF_1_ after freeze-thawing the sample (for full length gel see Fig. S3). (**C**) ATP hydrolysis activity of axle-less TF_1_ compared to the wild-type protein. Each data point represents one technical repeat. (**D**) Representative micrograph from the cryo-EM data collection. Scale bar represents 50 nm.

### 2.2 Overall structure of axle-less TF_1_

Single particle cryo-EM was used to determine the structure of the axle-less TF_1_. 1 mM MgAMP-PNP was added to the purified protein prior to cryo-EM grid freezing to ensure the appropriate nucleotide binding site occupancy of both the α and β subunits. Although the particles were distributed well on the cryo-EM grid, they exhibited preferred orientation (Fig. 1D) and so data was collected with stage tilt to reduce anisotropy [23]. During the refinement of the axle-less TF_1_ map, a CryoSPARC 3D classification resulted in a small subset of the total particles (2.2%) producing a much less anisotropic map where the density for the γ subunit was more complete. This could have been due to some flexibility of or damage to the γ subunit in most of the particles, but as no rotorless TF_1_ or alternate conformations of the axle-less γ subunit were observed, we postulate that this small set of particles may have rare orientations that allow better resolution of the γ subunit. The final map obtained from the 3D reconstructions contained density for all α, β and γ subunits (Fig. 2, S4A-B) allowing a molecular model to be built. The density for the foot of the γ subunit was not well resolved, consistent with other published single-particle cryo-EM TF_1_ maps, so this region was not completely modelled. Unexpectedly, the axle-less structure was most similar to the structure of the temperature sensitive TF_1_ in the binding-dwell conformation (PDB: 7L1Q, RMSD 1.5 Å, 2,869 residues), with the rotor foot in the same rotary position (Fig. S5). Although the original single-molecule rotation study performed on multiple axle-less TF_1_ mutations did not assign dwelling angles [21], we hypothesized that the axle-less ATPase would be observed in the catalytic-dwell state, because single-molecule rotation studies [13] have shown that F_1_-ATPases spend more time in the catalytic-dwell and this has been reflected in the majority of structural studies [4, 24]. In the binding-dwell state, γ is rotated counter-clockwise (when viewed from the membrane) by 40°, relative to its position in the catalytic-dwell state. This rotation has been suggested to be linked to the hydrolysis of ATP in one of the β subunits. As presented here, it appears that truncation of the γ axle led to TF_1_ adopting a binding-dwell rotatory position.

**Fig. 2.**
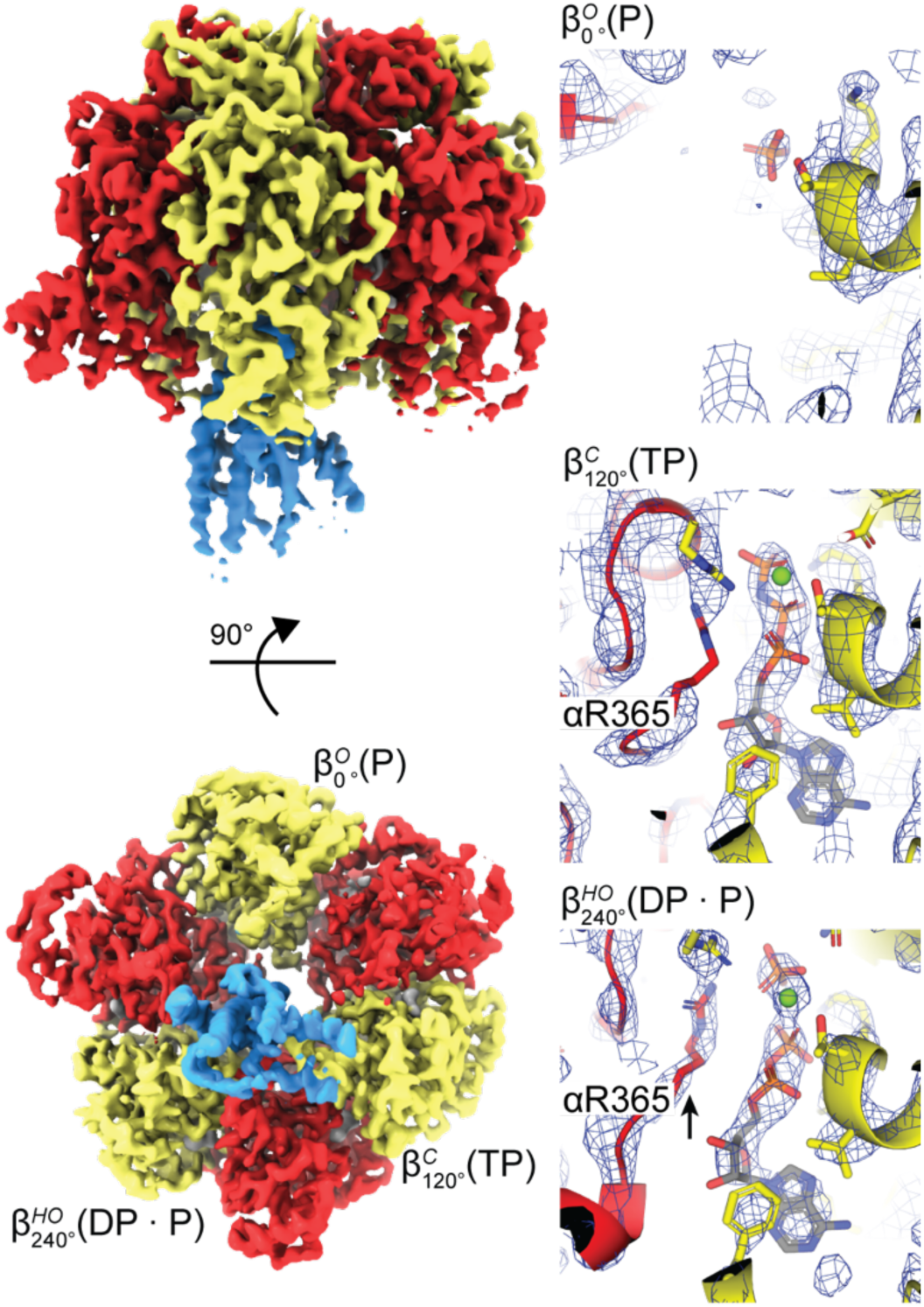
Structure of the axle-less TF_1_. The cryo-EM map (non-sharpened) is colored to show the α subunits in red, β subunits in yellow, γ in blue and bound nucleotides in white. The right-hand panels show the three β nucleotide binding sites, with the map (sharpened) overlaid as dark blue wire mesh 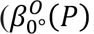; 3 σ 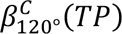 5 σ and 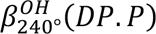; 7.5 σ). The β subunits are annotated using the nomenclature 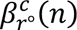 [11], where *c* is the conformation of the subunit (C = closed, HO = half-open and O = open), *r* is the approximate rotation angle in WT TF_1_ (as defined by single-molecule rotation studies [17, 25, 26]) and *n* is the nucleotide occupancy (TP = ATP or AMP-PNP, DP = ADP or AMP-PN, P = P_i_).

### 2.3 Conformation and nucleotide occupancies of the β subunits

All three β subunits in the axle-less TF_1_ structure adopt different conformational states with different nucleotide binding site occupancies (Fig. 2, Fig. 3). The open β subunit nucleotide binding site has density resembling an ion, and although SO_4_^2-^ has been observed at this location in related systems [27], we assigned this as P_i_ here because of the high concentration of phosphate in the purification buffer 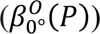. Interestingly, the phosphate back door seen in a previous study [11] is present in this open state (Fig. S7) providing further evidence that P_i_ may utilize a back door binding mechanism. The closed β subunit nucleotide binding site has density resembling a tri-phosphate nucleotide and so we assigned this as AMP-PNP because this had been added prior to imaging, although we refer to this state as TP for simplicity 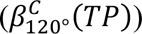. The half-open β subunit nucleotide binding site has density resembling ADP plus 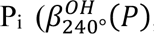, though given the limited resolution of these maps we cannot unequivocally attribute this density to ADP + P_i_. The presence of hydrolyzed products in this sample was unexpected, because AMP-PNP is often viewed as a competitive, nonhydrolyzable inhibitor of ATP-dependent enzymes [28]. It is possible that ADP may have been carried over from the purification steps, is a by-product of AMP-PNP degradation or partial hydrolysis by axle-less TF_1_ as has been observed in related systems [29]. Although the overall structure is similar to the temperature sensitive TF_1_ binding-dwell structure, the conformational state and nucleotide occupancy of one β subunit are different. The axle-less TF_1_ open state with P_i_ bound 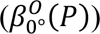 is not present in the TF_1_ binding-dwell structure and instead the equivalent subunit is half-closed and occupied by ATP 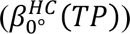. In the catalytic-dwell structure this β site is open but occupied by ADP; therefore, it could be hypothesized that the axle-less structure represents an intermediate step between the catalytic- and binding-dwell structures, where ADP has dissociated but ATP has not yet entered the binding site.

**Fig. 3.**
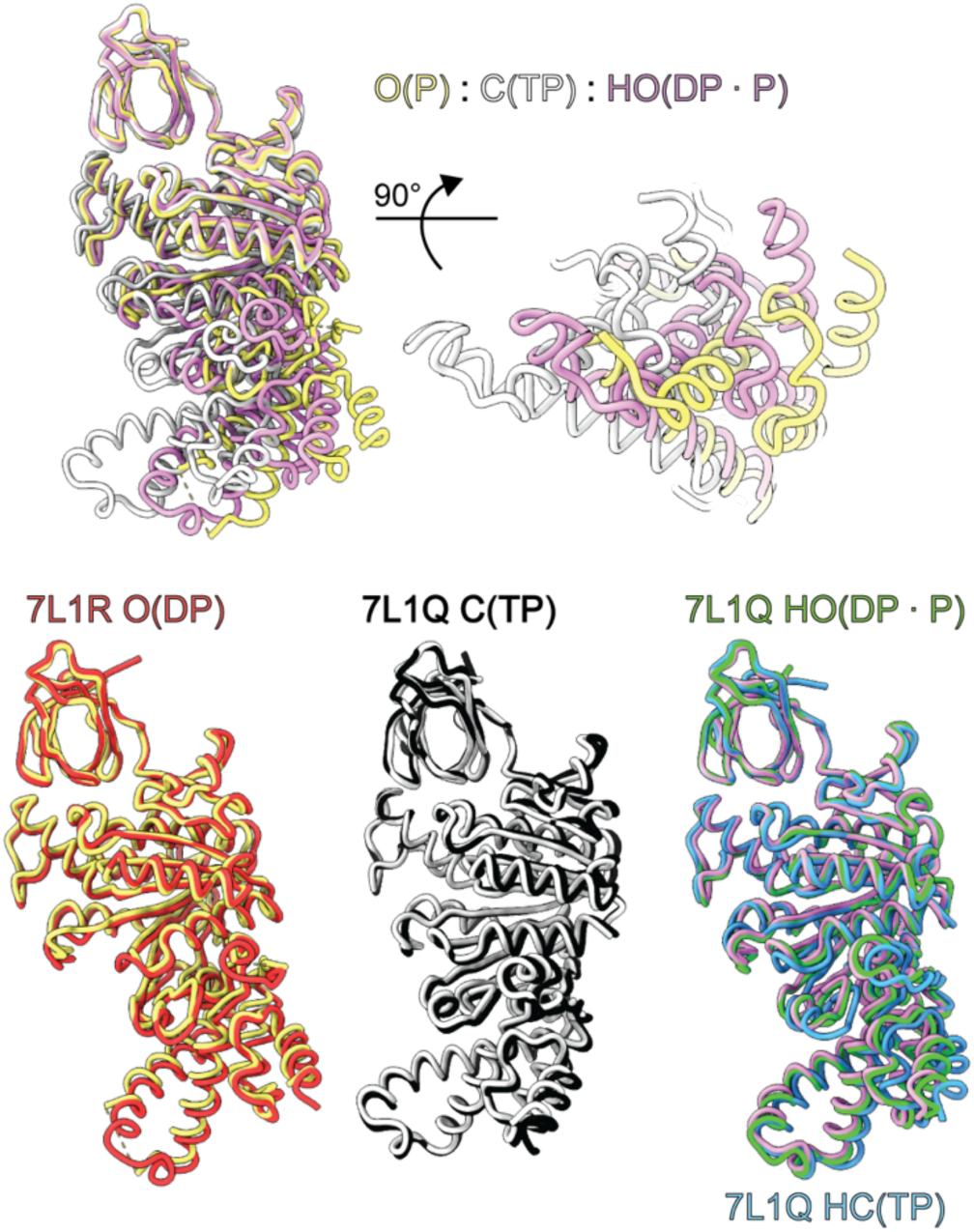
Conformations of the axle-less TF_1_ β subunits. Superpositions of the axle-less TF_1_ β subunits on each other and on β subunits from previous TF_1_ structures (PDB 7L1R and 7L1Q). Structures were aligned using the β crown structure (β1-82). The subunits are annotated based on their conformation and nucleotide occupancy; C = closed, HO = half-open, O = open, HC = half-closed and TP = ATP or AMP-PNP, DP = ADP or AMP-PN, P = P_i_.

### 2.4 Interactions of the axle-less γ subunit

There are three key sites where the β subunits interact with full-length γ in F_1_-ATPase: the hydrophobic sleeve, the β-catch loop, and the β-lever domains [4, 16, 17]. Given the γ N- and C-termini truncations in the axle-less F_1_-ATPase, the γ subunit is no longer able to interact with the hydrophobic sleeve or β-catch loop. However, enough of the axle remains to interact with the β-lever domains of the closed and half-open β subunits (Fig. 4). Residues 382-387 of the β^OH^ and β^C^ subunits contact the remaining axle of the γ subunit at regions γ15-26 and γ245-253 respectively (Fig. 4C). The β382-387 region includes the DIIA motif previously found to be important for F_1_ ATPase catalytic mechanism [16, 30, 31]. The γ foot remains in the same position as in the TF_1_ binding-dwell structure, indicating that the interactions between the β-lever domains and the remaining γ axle (Fig. 4) are strong enough the keep the γ subunit in place.

**Fig. 4.**
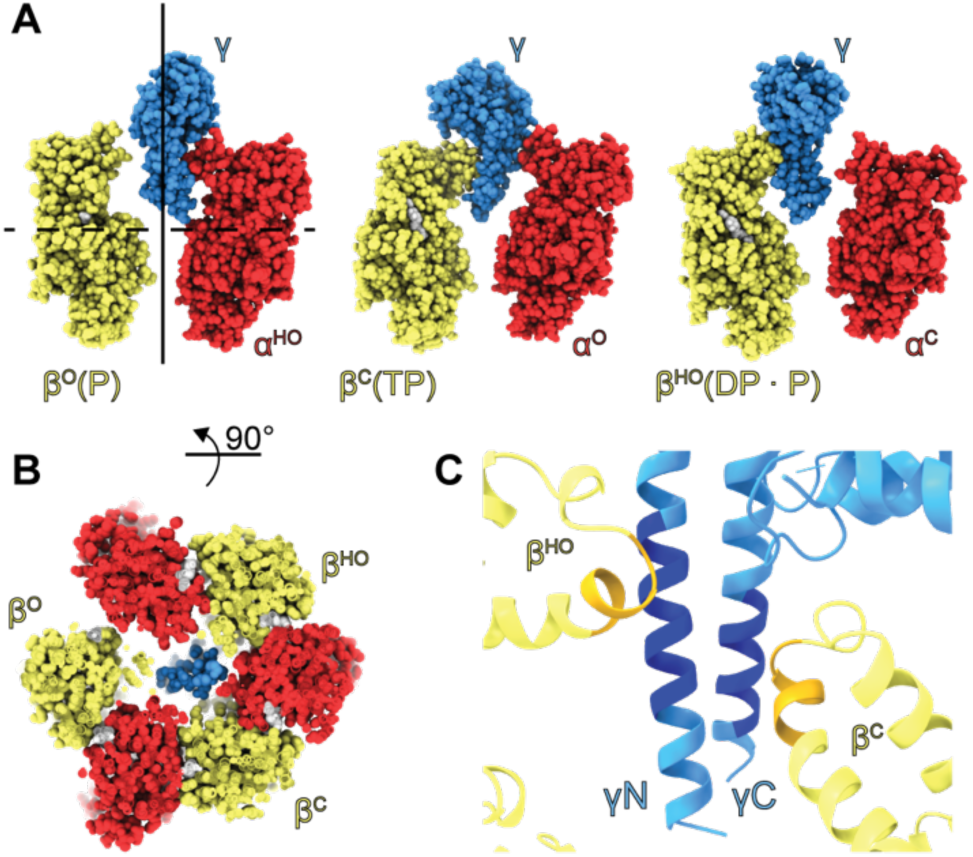
Interactions of axle-less γ with the β subunits. (**A)** Side views of the axle-less γ subunit with opposing αβ pairs. The solid black line indicates the γ rotation axis. Adapted from Furuike et al. [21]. (**B**) Cross-section through the axle-less TF_1_ structure. The plane of the cross-section is indicated by the dashed black line in (**A**). (**C**) A close-up of the interactions between the remaining γ-axle and the β-lever domains. The regions involved in the interactions (382-387 of the β^OH^ and β^C^ subunits and 15-26 and 245-253 of the γ subunit) are indicated by the darker colors. All panels are colored and β subunits are annotated as in Fig. 2.

Because the axle-less TF_1_ structure is in the binding-dwell rotary conformation, the interactions of γΔN4C25 with the β domains are different to those hypothesized by Furuike et al. [21]. In the axle-less TF_1_ structure, the β-lever domain of the open β subunit is disordered, meaning that this subunit has no contact with the γ subunit (Fig. 4A and B). This suggests that, without the axle, the γ subunit is unable to coordinate any conformational changes of the open β subunit when in the binding-dwell rotary position. Instead, the enzyme may rely on cooperativity from neighboring α and β subunits (as has been suggested in previous studies [32–34]) for ATP binding and closure of this subunit.

### 2.5 Structure of WT TF_1_ after addition of AMP-PNP

Because it was surprising that we observed axle-less TF_1_ in the binding-dwell rotary position, we performed single particle cryo-EM on wild-type (WT) TF_1_ protein after addition of 1 mM MgAMP-PNP (Fig. S3B, S4C-D, S8). This structure acted as a control to ensure that the rotary position we had observed with the mutant was due to the axle-less truncation, rather than the absence of the temperature sensitivity inducing βE190D mutation [11] or the addition of MgAMP-PNP vs MgATP. The map obtained of WT TF_1_ after the addition of 1 mM MgAMP-PNP is virtually identical to the temperature sensitive TF_1_ in the catalytic-dwell conformation (RMSD: 0.98 Å, 2,952 residues) except that nucleotide occupancy is different (Fig. S8D-E). In the temperature sensitive TF_1_ catalytic-dwell structure, ADP has clearly bound at 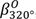, but in the WT TF_1_ +1 mM MgAMP-PNP structure 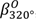 has only weak occupancy for an ion/water (Fig. S8D), suggesting the ADP observed in the temperature sensitive TF_1_ catalytic-dwell structure may be caused by the higher relative ADP concentration resulting from ATP hydrolysis in that study [11]. Because we observe WT TF_1_ +1 mM AMP-PNP in the catalytic-dwell, this suggests that the binding-dwell rotary position observed for axle-less TF_1_ in this study was due to the axle-less truncation.

### 2.6 Role of the γ subunit axle in efficient F_1_-ATPase catalysis

This study presents a structure of an axle-less TF_1_ that adopts the binding-dwell rotary conformation. Given that the binding-dwell state of the axle-less TF_1_ was captured, it is likely that the process of ATP binding and associated conformational changes are less efficient in this mutant, providing a possible explanation for the reduced catalytic activity of axle-less TF_1_. Previous studies have shown that the γ axle contacts three sites in the β subunit and disruption of these sites modulates F_1_-ATPase activity [16, 22, 35, 36]. The axle-less γ subunit used in this study is unable to interact with the hydrophobic sleeve and β-catch loop interaction sites and as a result, makes no contact with the β subunit that adopts an open conformation when the γ subunit is in the binding-dwell rotary conformation. Single molecule studies have also shown that the ATP binding rate is enhanced when γ is mechanically rotated in the direction of hydrolysis (i.e. counterclockwise when viewed from the membrane) [37]. Together these findings suggest that the interaction of the γ axle at these two sites is important for efficient ATP binding and coordinating the closure of open β subunits throughout the catalytic cycle. These interactions may enhance the ATP binding rate by stabilizing or guiding the conformational transition of the catalytic subunits, though the precise contribution and nature of each interaction requires further investigation. Overall, these findings highlight the importance of the complete γ subunit for efficient catalysis of TF_1_.

## 3. Materials and methods

### 3.1 Materials

Unless otherwise stated all materials and chemicals were obtained from Sigma-Aldrich.

### 3.2 Molecular biology

Nucleotides encoding a Strep-tag II (WSHPQFE) were inserted between the codons of residues N196 and K197 in the γ subunit of pHC95_γΔN4C25 (published previously [21]) by amplifying the plasmid with primers (Integrated DNA Technologies):

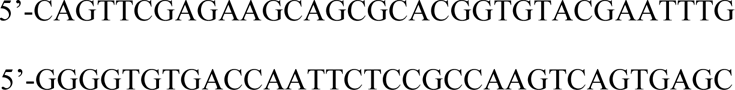

and Phusion DNA polymerase (New England Biolabs (NEB)), then treating the PCR product with KLD enzyme mix (NEB). The treated PCR product was transformed into *E. coli* JM109 competent cells (Promega) and the resulting colonies were cultured and mini-prepped (QIAGEN). The Strep-tag II insertion was confirmed by Sanger (Australian Genome Research Facility) and whole plasmid Nanopore sequencing (Plasmidsaurus, Oregon, US).

### 3.3 Protein expression and purification

Axle-less *Bacillus* sp. PS3 F_1_-ATPase was expressed in *E. coli* JM103Δ*unc* cells at 37°C for 19 h in 1 L of Terrific Broth with 100 µg/ml carbenicillin in a 2.5 L Tunair baffled flask, shaking at 180 rpm. No induction of expression was necessary. After expression, cells were harvested by centrifugation and stored at -20°C for future use. The cells were thawed and resuspended in 10 mL of Resuspension Buffer (50 mM Imidazole, 100 mM NaCl, pH 7) per 1 g of cell pellet. One cOmplete EDTA-free Protease Inhibitor Cocktail tablet (Roche) and a pinch of DNase I powder were added to the resuspended cell pellet before the cells were lysed by sonication (Branson digital sonifier, 30% power, 3 min) a 4°C. The lysate was then cleared by centrifugation at 42,000 × g for 50 min at 4°C before being applied to 2 mL of Ni-NTA resin (QIAGEN) in a gravity flow column, equilibrated with Resuspension Buffer. The column was washed with 3 x 10 mL of Resuspension Buffer before the bound protein was eluted with 2 x 5 mL of 500 mM Imidazole, 100 mM NaCl, pH 7 (IMAC Elution Buffer). The eluted sample was diluted with Resuspension Buffer to a final volume of 30 mL, before being applied to 2 mL of Strep-Tactin XT 4flow, High Capacity (IBA Lifesciences) resin in a gravity flow column, equilibrated with 1:2 IMAC Elution Buffer: Resuspension Buffer. The sample was applied twice to the Strep-Tactin column before being washed with 3 x 10 mL of Resuspension Buffer and eluted with 2 x 5mL of 50 mM Biotin in Resuspension Buffer. The protein was concentrated to 500 µL using a 100 kDa MWCO centrifugal concentrator (Amicon) before being injected onto a Superdex 200 10/300 GL column (Cytiva), equilibrated with 100 mM Potassium phosphate pH 7, 2 mM EDTA. Peak fractions were pooled, and flash frozen with liquid nitrogen before being stored at -80°C. Samples of each step of the purification were analysed by SDS-PAGE, using 4-12% Bolt Bis-Tris Plus Mini protein gels (Thermo Fisher/Invitrogen) stained with Quick Coomassie Stain (ProteinArk). Wild type *Bacillus* sp. PS3 F_1_-ATPase was expressed and purified as described above, but with the Strep-affinity purification step omitted.

### 3.4 ATPase activity assay

The ATPase activity of the F_1_-ATPase enzymes was assessed using the standard ATP regeneration assay [38]. This assay uses pyruvate kinase and lactate dehydrogenase to link the hydrolysis of ATP by the ATPase to the conversion of NADH to NAD^+^, which can be measured spectrophotometrically by the decrease in absorbance at 340 nm. Each reaction was performed in a 1 mL quartz cuvette at 298 K. The A_340_ of 5 units/ml of pyruvate kinase and lactate dehydrogenase, 1 mM ATP, 2 mM phosphor(enol)pyruvic acid (PEP), 100 mM potassium chloride, 50 mM MOPS, 2 mM magnesium chloride and 0.2 mM NADH was read, before the ATPase was quickly added and the decrease in absorbance measured over time. The final concentration of ATPase used the assay was 20 nM for the axle-less mutant and 20 nM, 5 nM or 2.5nM for the wild type. Multiple technical replicates were performed on three separate days. The first 20 seconds of the reaction was used to generate the rate of reaction, which was then converted to molecules of ATP hydrolysed by each ATPase enzyme per sec (s^-1^) using the extinction coefficient of NADH (ε340 = 6220 M^-1^cm^-1^) and the molar concentration of ATPase in the reaction.

### 3.5 Cryo-electron microscopy sample preparation

Axle-less TF_1_: To ensure the nucleotide occupancy of all binding sites, 6 mg/ml (Abs 0.1% = 0.41) Axle-less PS3 F_1_-ATPase was incubated with 1 mM MgAMP-PNP for 20 min. 3.5 µL of this sample was then applied to UltrAuFoil R1.2/1.3, 300 mesh holey gold supports (Quantifoil), which had been glow discharged at 15 mA for 1 min (PELCO easiGlow). The grids were blotted for 4 s at 20°C, 100 % humidity, with blot force set to 0 and plunge frozen into liquid ethane using a Vitrobot Mark IV (FEI).

WT TF_1_: 4.2 mg/ml (Abs 0.1% = 0.39) WT PS3 F_1_-ATPase was incubated with 1 mM MgAMP-PNP. 3.5 µL of this sample was then applied to UltrAuFoil R1.2/1.3, 300 mesh holey gold supports (Quantifoil), which had been glow discharged at 15 mA for 1 min (PELCO easiGlow). The grids were blotted for 4 s at 20°C, 100 % humidity, with blot force set to 0 and plunge frozen into liquid ethane using a Vitrobot Mark IV (FEI). Cryo-EM data collection summarized in Table S1.

### 3.6 Cryo-EM data collection and processing

Axle-less TF_1_: The cryo-EM samples were screened for ice thickness and particle density using a Talos Arctica (Thermo Fisher Scientific) transmission electron microscope (TEM), operating at 200 kV. For data collection, the screened grids were transferred to a Titan Krios (Thermo Fisher Scientific) TEM operating at 300 kV, equipped with a BioQuantum energy filter (Gatan) (15 eV) and a K3 direct detection camera (Gatan). To overcome the orientation bias of the particles, data were collected with stage tilt. Approximately two thirds of the data were collected at 30°, with the remaining third split over 0°, 10° and 20°. Data were collected using EPU (E Pluribus Unum - Thermo Fisher Scientific) at 60,000X magnification (165,000X was displayed on user interface due to the energy filter), resulting in a pixel size of 0.82 Å. A defocus range of 0.5-1.5 nm was used. A total dose of 59 e^-^ Å^-2^ with an exposure time of 5 s was applied over 75 frames. A total of 2,992 exposures were collected.

The data were processed using cryoSPARC (v3.3.2+220824). After motion correction and CTF estimation, poor quality exposures were discarded based on CTF fit, motion and ice thickness. The particles were then auto picked with the circular blob picker (min diameter 100 Å, max diameter 200 Å) and extracted with a box size of 250 px. Multiple rounds of 2D classification were used to clean up the particle selection, resulting in 1,056,364 particles. An *ab initio* 3D model was generated and improved with subsequent rounds of homogeneous refinement and non-uniform refinement. At this stage, the map was still highly anisotropic, but a round of 3D classification (as implemented in cryoSPARC v4.0.1) with “force hard classification” turned on into 2 classes, resulted in a greatly improved map with only small subset of the total particles (23,813). This map was further refined using homogeneous refinement and non-uniform refinement, before being used for model building.

WT TF_1_: The cryo-EM samples were screened for ice thickness and particle density using a Talos Arctica (Thermo Fisher Scientific) transmission electron microscope (TEM), operating at 200 kV. Data was collected on a Talos Arctica (Thermo Fisher Scientific) TEM operating at 200 kV, equipped with a Falcon III camera (Thermo Fisher Scientific) operating in counting mode. To overcome the orientation bias of the particles, all data were collected with 30° stage tilt. Data were collected using EPU (Thermo Fisher Scientific) at 150,000X magnification, resulting in a pixel size of 0.986 Å. A defocus range of -1.5 to -2.5 nm was used. A total dose of 50 e^-^ Å^-2^ with an exposure time of 55 s was applied over 60 frames. A total of 1,467 exposures were collected.

The data were processed using cryoSPARC (v4.4.1). After motion correction and CTF estimation, poor quality exposures were discarded based on CTF fit, motion and ice thickness. The particles were then auto picked with the circular blob picker (min diameter 100 Å, max diameter 150 Å) and extracted with a box size of 256 px. All further processing steps were performed in cryoSPARC. Multiple rounds of 2D classification were used to clean up the particle selection, resulting in 666,517 particles. An *ab initio* 3D model was generated and improved with subsequent rounds of homogeneous refinement and non-uniform refinement. The map was still highly anisotropic, but 3D classification (as implemented in cryoSPARC v4.4.1) with forced classification into 10 classes, resulted in an improved map. However, the resultant map was still anisotropic and therefore cryoFEM [39] was used to sharpen the map and aid in interpretation.

Cryo-EM data processing summarized in Table S1.

### 3.7 Model building and structure refinement

Axle-less TF_1_: The initial model of the axle-less F_1_-ATPase was built using ModelAngelo (v0.1) [40], with the sharpened map and sequence as input. Several rounds of manual adjustment in Coot, using the published binding-dwell structure (PDB: 7L1Q) as a guide, and real-space refinement in Phenix were performed, with reference to the MolProbity validation statistics. ISOLDE, as implemented in ChimeraX-1.5, was used throughout refinement to improve the geometry of the model.

WT TF_1_: The E190D TF_1_ catalytic dwell structure (PDB: 7L1R [11]) was docked into the map using Chimera1.5 [41]. Position β190 is mutated to E (the WT sequence) and the model was iteratively built in Coot [42] and real-space refined in Phenix [43] and ISOLDE [44].

All structure figures were made using ChimeraX-1.5 or PyMOL. RMSD calculations were performed using Secondary Structure Matching in Coot.

Model statistics summarized in Table S1.

## CRediT author contributions

Emily J. Furlong: Methodology, Investigation, Formal analysis, Visualisation, Writing - Original Draft. Ian B. P. Reiniger-Chatzigian: Investigation. Yi C. Zeng: Investigation. Simon H. J. Brown: Investigation. Meghna Sobti: Investigation. Alastair G. Stewart: Conceptualization, Funding acquisition, Formal analysis, Visualisation, Writing - Review & Editing.

## Data availability

The axle-less TF_1_ model and maps have been deposited under the Protein Data Bank (PDB) code 8U1H and Electron Microscopy Data Bank (EMDB) code EMD-41811, respectively. The axle-less cryo-EM raw data was deposited under the Electron Microscopy Public Image Archive [45] code EMPIAR-12183. The wild-type TF_1_ model and maps have been deposited under PDB 9AVJ and EMD-43903, respectively.

## Acknowledgements

We wish to thank Shou Furuike (Osaka Medical and Pharmaceutical University) for the axle-less TF _1_ expression plasmid, as well as Hiroyuki Noji and Hiroshi Ueno (University of Tokyo) for critically reviewing the manuscript. We also thank and acknowledge the use of the University of Wollongong Cryogenic Electron Microscopy Facility at Molecular Horizons, as well as the use of the Victor Chang Innovation Centre (funded by the NSW Government) and the Electron Microscope Unit at UNSW Sydney. Molecular graphics and analyses performed with UCSF ChimeraX, developed by the Resource for Biocomputing, Visualization, and Informatics at the University of California, San Francisco, with support from National Institutes of Health R01-GM129325 and the Office of Cyber Infrastructure and Computational Biology, National Institute of Allergy and Infectious Diseases. A.G.S. was supported by a National Health and Medical Research Council APP2016308 and APP1146403. E.J.F. was supported by an Australian National University (ANU) Futures Award. This research was conducted by the Australian Research Council Industrial Transformation Training Centre for Cryo-Electron Microscopy of Membrane Proteins for Drug Discovery (IC200100052).

## SUPPLEMENTARY DATA

**Figure S1:**
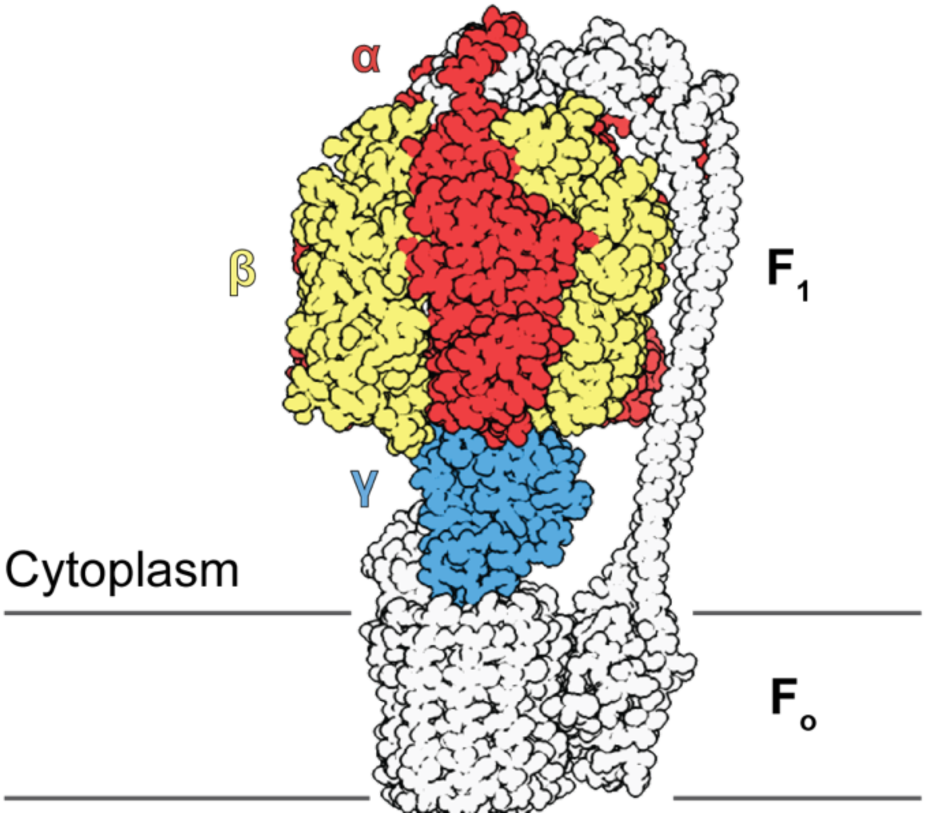
Structure of F_1_F_o_ ATP synthase. Molecular model of the *Geobacillus stearothermophilus* (also known as *Bacillus sp.* PS3) F_1_F_o_ ATP synthase (PDB: 6N2Y). The core F_1_ (α, β, γ) components are coloured, and the rest of the model is displayed in white.

**Figure S2:**
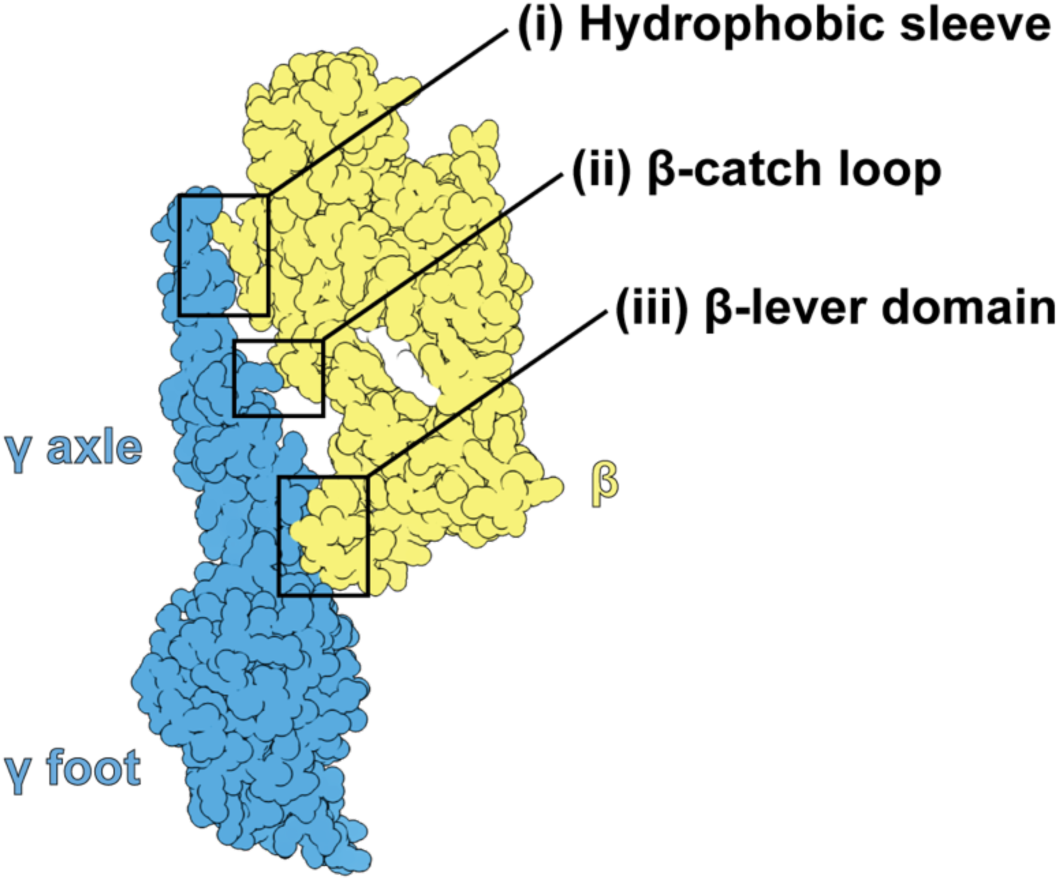
Contact sites between the γ and β subunits in F_1_ ATPase. The β subunit is shown in yellow and the γ subunit in blue. The three key interaction sites are (i) the hydrophobic sleeve, (ii) the β-catch loop and (iii) the β-lever domain and these contact the γ axle.

**Figure S3:**
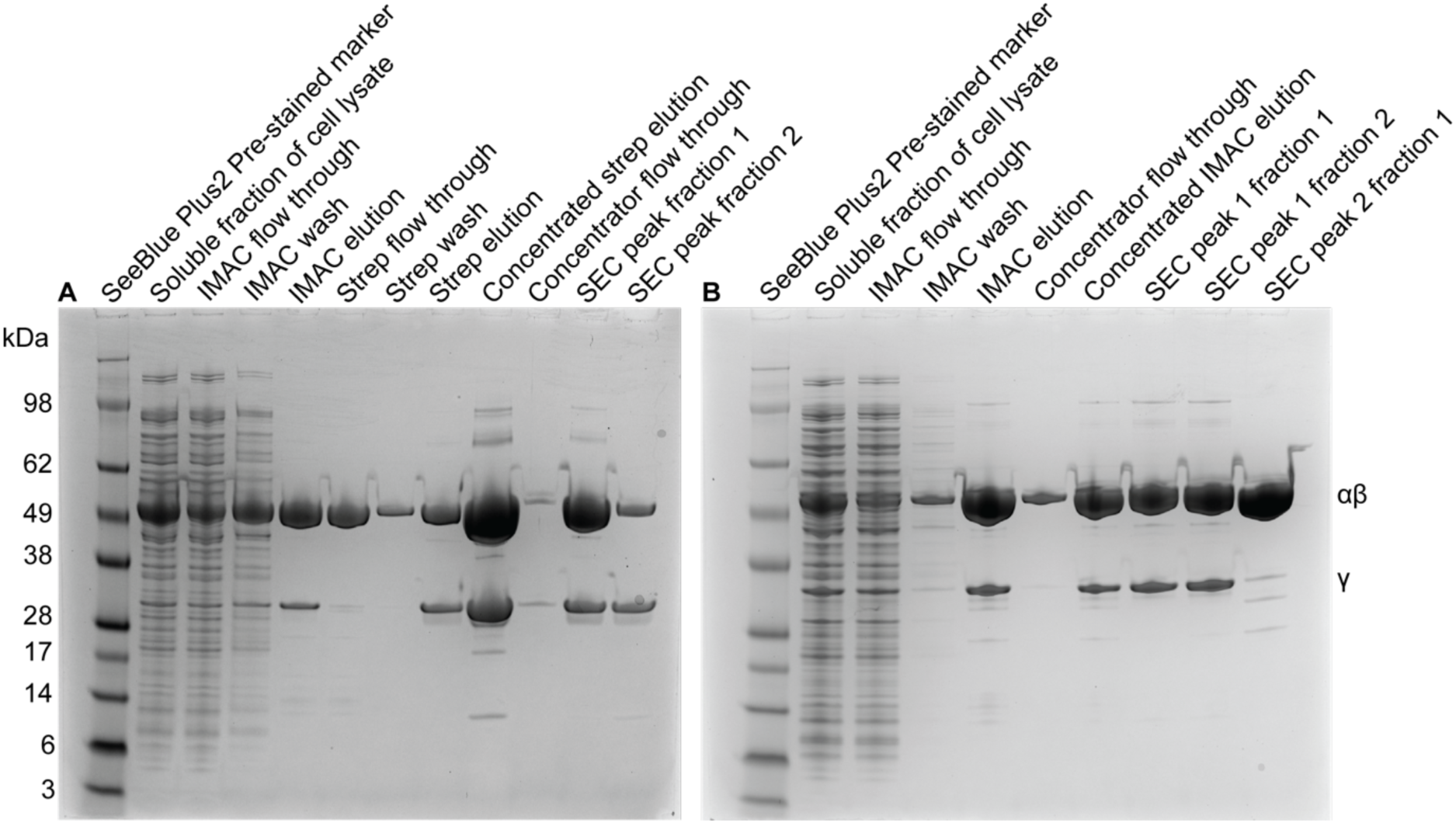
SDS-PAGE from protein purifications. (**A**) SDS-PAGE of the samples taken throughout the axle-less TF_1_ protein purification. (**B**) SDS-PAGE of the samples taken throughout the wild-type TF_1_ protein purification.

**Figure S4:**
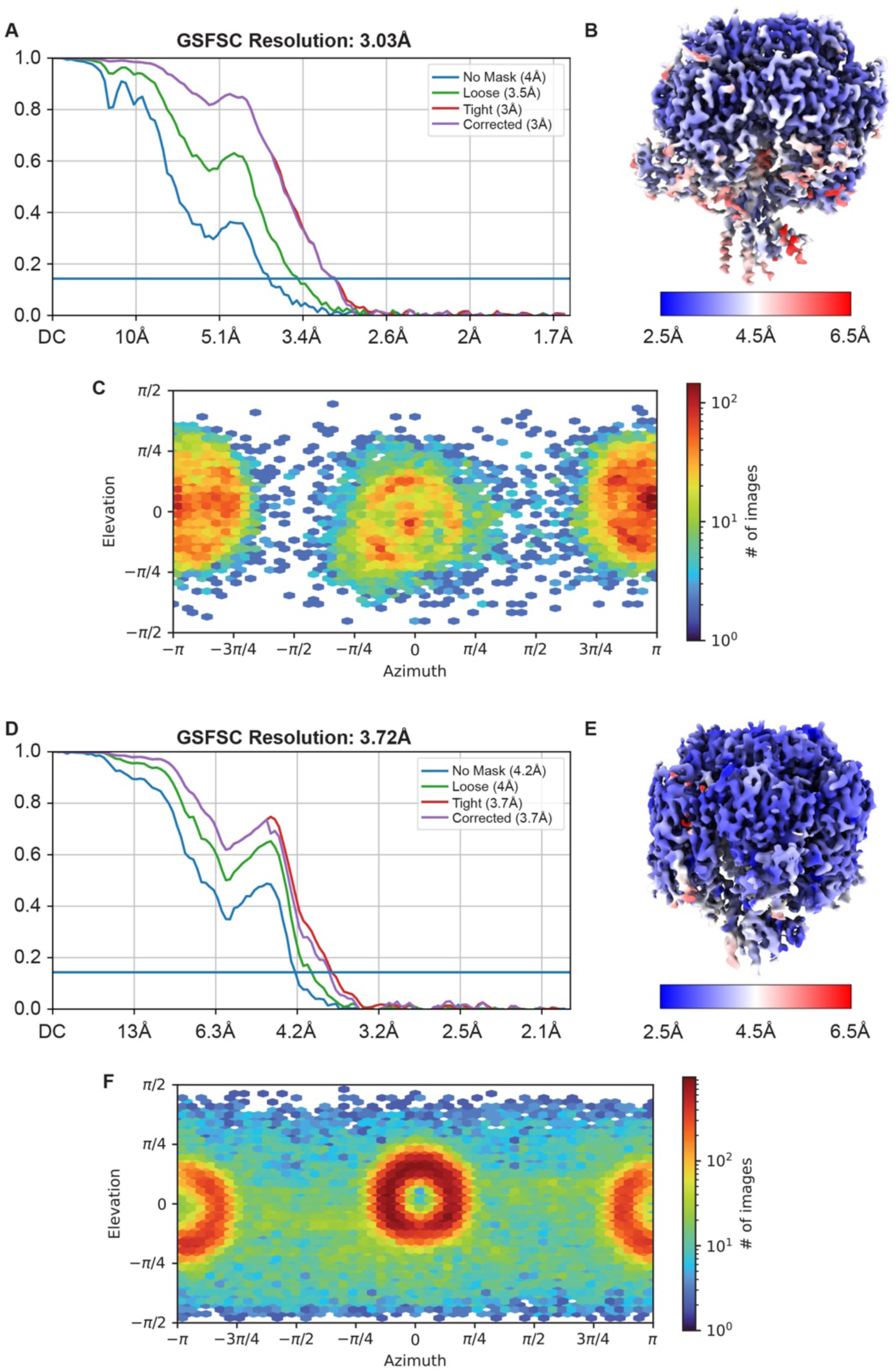
Resolution estimates for cryo-EM maps. Gold standard Fourier shell correlation (GSFSC) curves from cryoSPARC for the (**A**) axle-less TF_1_ map and (**D)** wild-type TF_1_ map. Sharpened (**B**) axle-less TF_1_ map and (**E**) wild-type map coloured by local resolution estimate calculated in cryoSPARC. Viewing direction distribution plot (**C**) axle-less TF_1_ map and (**F)** wild-type map.

**Figure S5:**
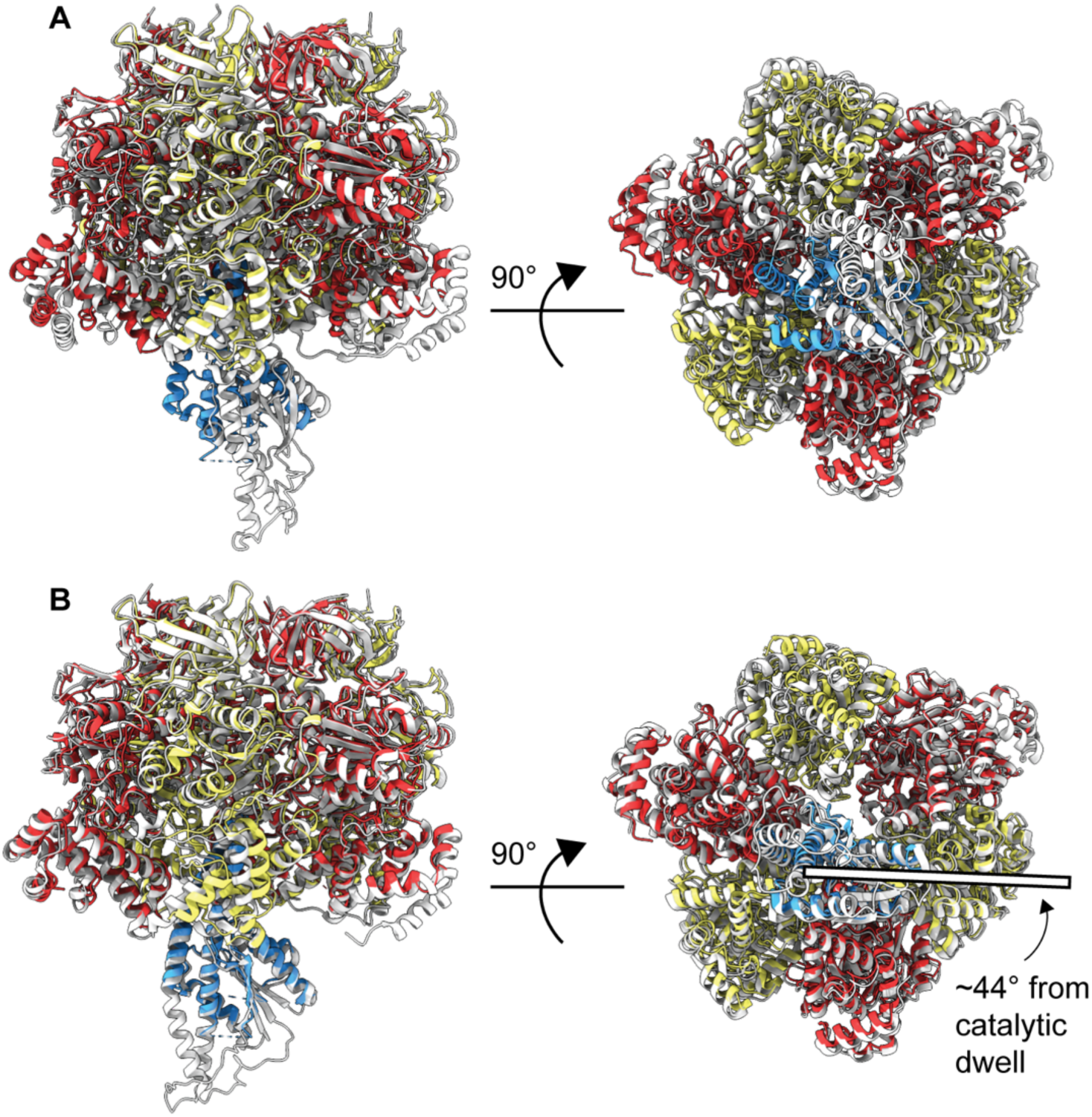
Comparison of the axle-less TF_1_ structure to other TF_1_ structures. The axle-less structure (colored) aligned to the (**A**) TF_1_ catalytic-dwell state (PDB: 7L1R, grey; RMSD 2.2 Å, 2,624 residues) and (**B**) TF_1_ binding-dwell state (PDB: 7L1Q, grey; RMSD 1.5 Å, 2,869 residues). For the figure, the structures were aligned using the crown region (residues 1-82) of the α and β subunits.

**Figure S6:**
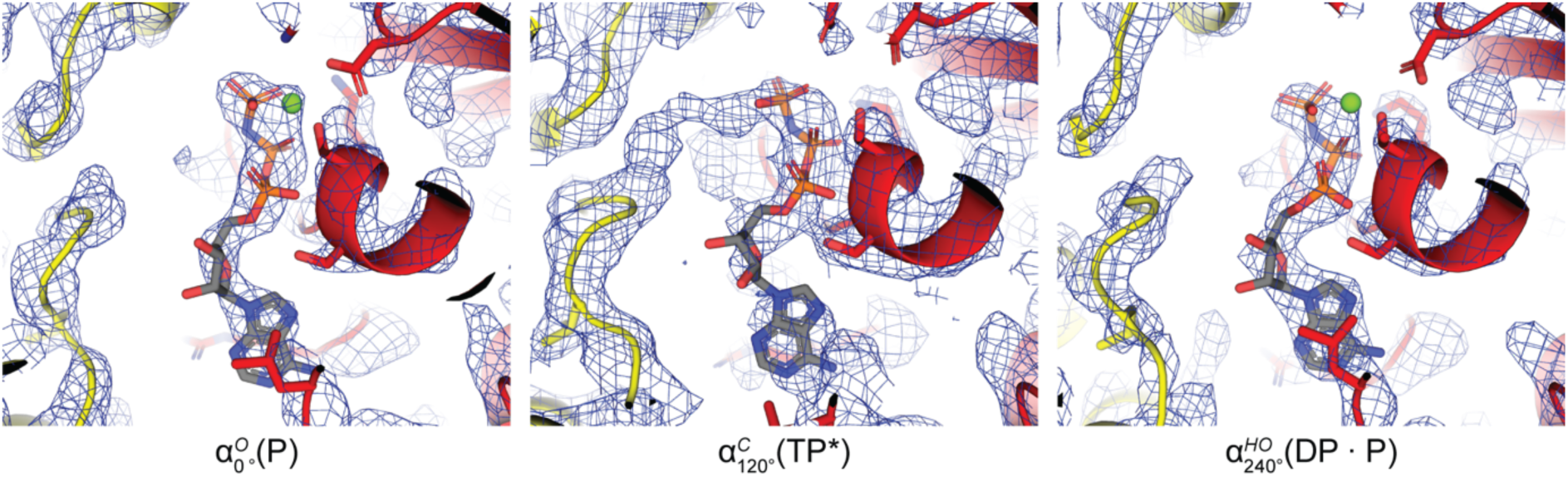
Occupancy of the ATP binding sites in the α subunits of the axle-less TF_1_ structure. The map is shown as dark blue wire mesh 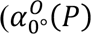; 5 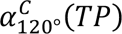; 3.1 σ and 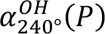.; 6 σ,

**Figure S7:**
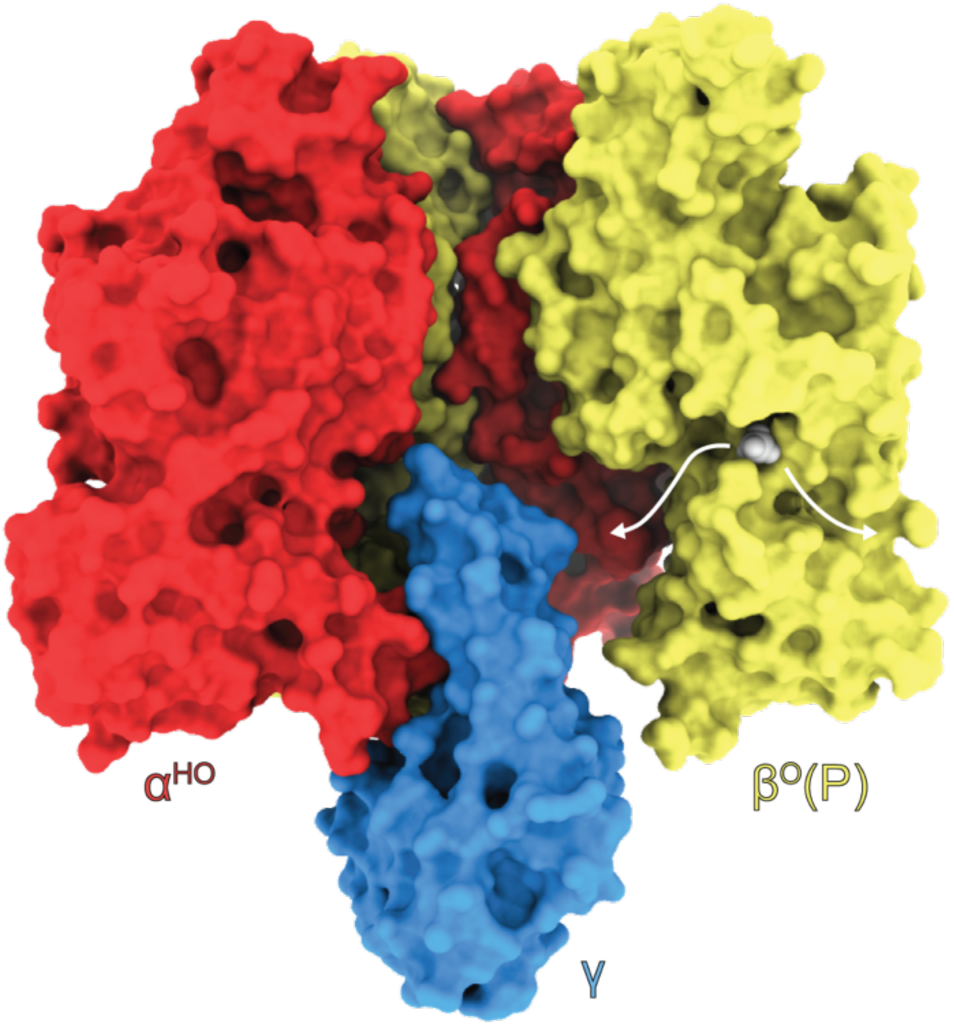
Inorganic phosphate tunnel in the axle-less TF_1_ structure. The axle-less structure is displayed in surface representation and coloured as in Fig 2. One α and one β subunit have been removed to reveal the inorganic phosphate tunnel, which is indicated by the white arrows.

**Figure S8:**
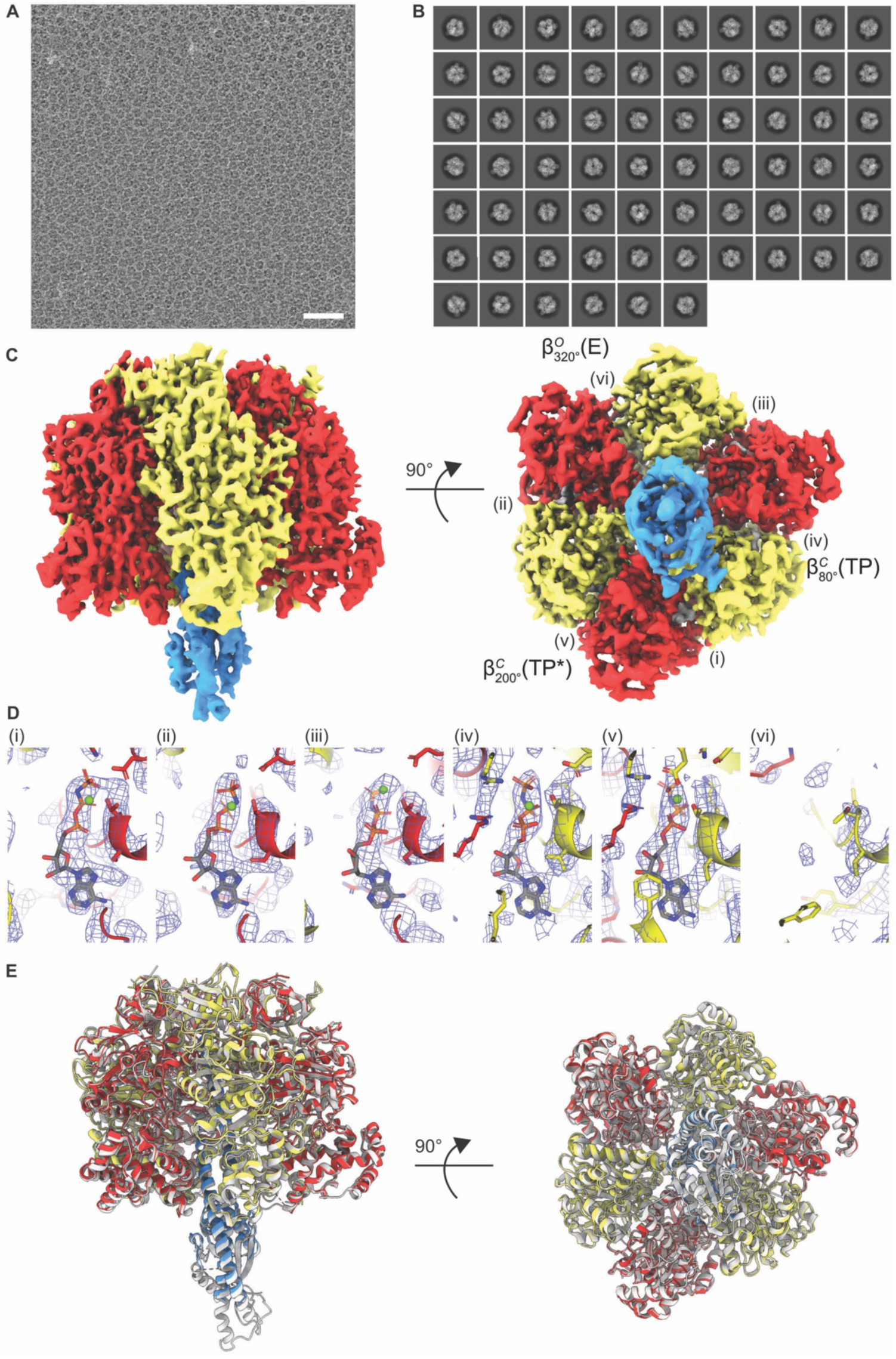
Wild-type (WT) TF_1_ structure. (**A**) Representative micrograph from the WT TF_1_ cryo-EM data collection. Scale bar represents 50 nm. (**B**) 2D classes from the WT TF_1_ data processing. (**C**) The cryoFEM map of the WT TF_1_ ATPase is colored to show the α subunits in red, β subunits in yellow, γ in blue and bound nucleotides in white. (**D**) The nucleotide binding sites of the α and β subunits, with the CryoSPARC sharpened map overlaid as dark blue wire mesh (6 σ in all panels). (**E**) The WT TF_1_ structure (colored) aligned to the TF_1_ catalytic dwell state (PDB: 7L1R, grey; RMSD 0.98 Å, 2,952 residues). For the figure, the structures were aligned using the crown region (residues 1-82) of the α and β subunits.

**Supplementary Table 1:** Cryo-EM data collection, refinement, and validation statistics for the axle-less TF_1_ and wild-type TF_1_ structures.

|  | Axle-less TF<sub>1</sub> AMPPNP<br>(EMD-41811)<br>(PDB 8U1H) | Wild-type TF<sub>1</sub><br>AMPPNP<br>(EMD-43903)<br>(PDB 9AVJ) |
| --- | --- | --- |
| **Data collection and processing** |  |  |
| Magnification | 60,000X* | 150,000X |
| Voltage (kV) | 300 | 200 |
| Electron exposure (e-/Å<sup>2</sup>) | 59 | 50 |
| Defocus range (μm) | 0.5-1.5 | 1.5-2.5 |
| Pixel size (Å) | 0.82 | 0.99 |
| Symmetry imposed | C1 | C1 |
| Initial particle images (no.) | 1,555,897 | 1,022,877 |
| Final particle images (no.) | 23,813 | 153,828 |
| Map resolution (Å) | 3.03 | 3.72 |
| FSC threshold | 0.143 | 0.143 |
| **Refinement** |  |  |
| Initial model used (PDB code) | NA | 7L1R |
| Model resolution (Å) | 3.3 | 4.3 |
| FSC threshold | 0.5 | 0.5 |
| Map sharpening B factor (Å<sup>2</sup>) | 51.2 | 134.1 |
| Model composition |  |  |
| Non-hydrogen atoms | 22,720 | 23,188 |
| Protein residues | 2,926 | 2988 |
| Ligands | 11 | 10 |
| B factors (Å<sup>2</sup>) |  |  |
| Protein | 65 | 44 |
| Ligand | 70 | 46 |
| R.m.s. deviations |  |  |
| Bond lengths (Å) | 0.004 | 0.009 |
| Bond angles (°) | 0.600 | 1.017 |
| Validation |  |  |
| MolProbity score | 1.9 | 1.7 |
| Clashscore | 10 | 9 |
| Poor rotamers (%) | 0.5 | 0.45 |
| Ramachandran plot |  |  |
| Favored (%) | 95.23 | 96.58 |
| Allowed (%) | 4.77 | 3.38 |
| Disallowed (%) | 0.00 | 0.03 |

\*165,000X on user interface Mean B factors

## References

[1] J.E. Walker, The ATP synthase: the understood, the uncertain and the unknown, Biochem Soc Trans, 41 (2013) 1–16.

[2] W. Kuhlbrandt, Structure and Mechanisms of F-Type ATP Synthases, Annu Rev Biochem, (2019).

[3] A.G. Stewart, E.M. Laming, M. Sobti, D. Stock, Rotary ATPases - dynamic molecular machines, Curr Opin Struct Biol, 25 (2014) 40–48.

[4] J.P. Abrahams, A.G. Leslie, R. Lutter, J.E. Walker, Structure at 2.8 Å resolution of F_1_-ATPase from bovine heart mitochondria, Nature, 370 (1994) 621–628.

[5] H. Noji, R. Yasuda, M. Yoshida, K. Kinosita, Jr., Direct observation of the rotation of F_1_-ATPase, Nature, 386 (1997) 299–302.

[6] P.D. Boyer, The ATP synthase - a splendid molecular machine, Annu Rev Biochem, 66 (1997) 717–749.

[7] T.M. Duncan, V.V. Bulygin, Y. Zhou, M.L. Hutcheon, R.L. Cross, Rotation of subunits during catalysis by Escherichia coli F_1_-ATPase, Proc Natl Acad Sci U S A, 92 (1995) 10964–10968.

[8] M. Yoshida, N. Sone, H. Hirata, Y. Kagawa, A highly stable adenosine triphosphatase from a thermophillie bacterium. Purification, properties, and reconstitution, J Biol Chem, 250 (1975) 7910–7916.

[9] Y. Shirakihara, A.G. Leslie, J.P. Abrahams, J.E. Walker, T. Ueda, Y. Sekimoto, M. Kambara, K. Saika, Y. Kagawa, M. Yoshida, The crystal structure of the nucleotide-free alpha 3 beta 3 subcomplex of F_1_-ATPase from the thermophilic *Bacillus* PS3 is a symmetric trimer, Structure, 5 (1997) 825–836.

[10] R. Yasuda, H. Noji, K. Kinosita, Jr., M. Yoshida, F_1_-ATPase is a highly efficient molecular motor that rotates with discrete 120 degree steps, Cell, 93 (1998) 1117–1124.

[11] M. Sobti, H. Ueno, H. Noji, A.G. Stewart, The six steps of the complete F_1_-ATPase rotary catalytic cycle, Nat Commun, 12 (2021) 4690.

[12] S. Enoki, R. Watanabe, R. Iino, H. Noji, Single-molecule study on the temperature-sensitive reaction of F_1_-ATPase with a hybrid F_1_ carrying a single beta(E190D), J Biol Chem, 284 (2009) 23169–23176.

[13] K. Adachi, R. Yasuda, H. Noji, H. Itoh, Y. Harada, M. Yoshida, K. Kinosita, Jr., Stepping rotation of F_1_-ATPase visualized through angle-resolved single-fluorophore imaging, Proc Natl Acad Sci U S A, 97 (2000) 7243–7247.

[14] R. Watanabe, R. Iino, K. Shimabukuro, M. Yoshida, H. Noji, Temperature-sensitive reaction intermediate of F_1_-ATPase, EMBO Rep, 9 (2008) 84–90.

[15] T. Elston, H. Wang, G. Oster, Energy transduction in ATP synthase, Nature, 391 (1998) 510–513.

[16] W.D. Frasch, Z.A. Bukhari, S. Yanagisawa, F_1_FO ATP synthase molecular motor mechanisms, Front Microbiol, 13 (2022) 965620.

[17] J.L. Martin, R. Ishmukhametov, T. Hornung, Z. Ahmad, W.D. Frasch, Anatomy of F_1_-ATPase powered rotation, Proc Natl Acad Sci U S A, 111 (2014) 3715–3720.

[18] T. Uchihashi, R. Iino, T. Ando, H. Noji, High-speed atomic force microscopy reveals rotary catalysis of rotorless F(1)-ATPase, Science, 333 (2011) 755–758.

[19] M. Sobti, H. Ueno, S.H.J. Brown, H. Noji, A.G. Stewart, The series of conformational states adopted by rotorless F(1)-ATPase during its hydrolysis cycle, Structure, (2024).

[20] M.D. Hossain, S. Furuike, Y. Maki, K. Adachi, T. Suzuki, A. Kohori, H. Itoh, M. Yoshida, K. Kinosita, Jr., Neither helix in the coiled coil region of the axle of F_1_-ATPase plays a significant role in torque production, Biophys J, 95 (2008) 4837–4844.

[21] S. Furuike, M.D. Hossain, Y. Maki, K. Adachi, T. Suzuki, A. Kohori, H. Itoh, M. Yoshida, K. Kinosita, Jr., Axle-less F_1_-ATPase rotates in the correct direction, Science, 319 (2008) 955–958.

[22] K.W. Boltz, W.D. Frasch, Hydrogen bonds between the alpha and beta subunits of the F_1_-ATPase allow communication between the catalytic site and the interface of the beta catch loop and the gamma subunit, Biochemistry, 45 (2006) 11190–11199.

[23] K. Naydenova, C.J. Russo, Measuring the effects of particle orientation to improve the efficiency of electron cryomicroscopy, Nat Commun, 8 (2017) 629.

[24] M. Sobti, J.L. Walshe, D. Wu, R. Ishmukhametov, Y.C. Zeng, C.V. Robinson, R.M. Berry, A.G. Stewart, Cryo-EM structures provide insight into how *E. coli* F_1_F_o_ ATP synthase accommodates symmetry mismatch, Nat Commun, 11 (2020) 2615.

[25] R. Yasuda, H. Noji, M. Yoshida, K. Kinosita, Jr., H. Itoh, Resolution of distinct rotational substeps by submillisecond kinetic analysis of F_1_-ATPase, Nature, 410 (2001) 898–904.

[26] T. Bilyard, M. Nakanishi-Matsui, B.C. Steel, T. Pilizota, A.L. Nord, H. Hosokawa, M. Futai, R.M. Berry, High-resolution single-molecule characterization of the enzymatic states in Escherichia coli F_1_-ATPase, Philos Trans R Soc Lond B Biol Sci, 368 (2013) 20120023.

[27] G. Cingolani, T.M. Duncan, Structure of the ATP synthase catalytic complex F_1_ from *Escherichia coli* in an autoinhibited conformation, Nat Struct Mol Biol, 18 (2011) 701–707.

[28] H.S. Penefsky, Differential effects of adenylyl imidodiphosphate on adenosine triphosphate synthesis and the partial reactions of oxidative phosphorylation, J Biol Chem, 249 (1974) 3579–3585.

[29] Y. Suzuki, T. Shimizu, H. Morii, M. Tanokura, Hydrolysis of AMPPNP by the motor domain of ncd, a kinesin-related protein, FEBS Lett, 409 (1997) 29–32.

[30] E. Usukura, T. Suzuki, S. Furuike, N. Soga, E. Saita, T. Hisabori, K. Kinosita, Jr., M. Yoshida, Torque generation and utilization in motor enzyme F0F_1_-ATP synthase: half-torque F_1_ with short-sized pushrod helix and reduced ATP Synthesis by half-torque F0F_1_, J Biol Chem, 287 (2012) 1884–1891.

[31] D.S. Lowry, W.D. Frasch, Interactions between beta D372 and gamma subunit N-terminus residues gamma K9 and gamma S12 are important to catalytic activity catalyzed by Escherichia coli F_1_F0-ATP synthase, Biochemistry, 44 (2005) 7275–7281.

[32] L. Dai, H. Flechsig, J. Yu, Deciphering Intrinsic Inter-subunit Couplings that Lead to Sequential Hydrolysis of F(1)-ATPase Ring, Biophys J, 113 (2017) 1440–1453.

[33] P. Falson, A. Goffeau, M. Boutry, J.M. Jault, Structural insight into the cooperativity between catalytic and noncatalytic sites of F_1_-ATPase, Biochim Biophys Acta, 1658 (2004) 133–140.

[34] K. Braig, R.I. Menz, M.G. Montgomery, A.G. Leslie, J.E. Walker, Structure of bovine mitochondrial F(1)-ATPase inhibited by Mg(2+) ADP and aluminium fluoride, Structure, 8 (2000) 567–573.

[35] K.W. Boltz, W.D. Frasch, Interactions of gamma T273 and gamma E275 with the beta subunit PSAV segment that links the gamma subunit to the catalytic site Walker homology B aspartate are important to the function of Escherichia coli F_1_F0 ATP synthase, Biochemistry, 44 (2005) 9497–9506.

[36] M.D. Greene, W.D. Frasch, Interactions among gamma R268, gamma Q269, and the beta subunit catch loop of Escherichia coli F_1_-ATPase are important for catalytic activity, J Biol Chem, 278 (2003) 51594–51598.

[37] R. Watanabe, D. Okuno, S. Sakakihara, K. Shimabukuro, R. Iino, M. Yoshida, H. Noji, Mechanical modulation of catalytic power on F_1_-ATPase, Nat Chem Biol, 8 (2011) 86–92.

[38] M. Sobti, R. Ishmukhametov, A.G. Stewart, ATP Synthase: Expression, Purification, and Function, Methods Mol Biol, 2073 (2020) 73–84.

[39] X. Dai, L. Wu, S. Yoo, Q. Liu, Integrating AlphaFold and deep learning for atomistic interpretation of cryo-EM maps, bioRxiv, (2023) 2023.2002.2002.526877.

[40] K. Jamali, L. Kall, R. Zhang, A. Brown, D. Kimanius, S.H.W. Scheres, Automated model building and protein identification in cryo-EM maps, Nature, (2024).

[41] T.D. Goddard, C.C. Huang, E.C. Meng, E.F. Pettersen, G.S. Couch, J.H. Morris, T.E. Ferrin, UCSF ChimeraX: Meeting modern challenges in visualization and analysis, Protein Sci, 27 (2018) 14–25.

[42] P. Emsley, B. Lohkamp, W.G. Scott, K. Cowtan, Features and development of Coot, Acta Crystallogr D Biol Crystallogr, 66 (2010) 486–501.

[43] P.V. Afonine, B.K. Poon, R.J. Read, O.V. Sobolev, T.C. Terwilliger, A. Urzhumtsev, P.D. Adams, Real-space refinement in PHENIX for cryo-EM and crystallography, Acta Crystallogr D Struct Biol, 74 (2018) 531–544.

[44] T.I. Croll, ISOLDE: a physically realistic environment for model building into low-resolution electron-density maps, Acta Crystallogr D Struct Biol, 74 (2018) 519–530.

[45] A. Iudin, P.K. Korir, S. Somasundharam, S. Weyand, C. Cattavitello, N. Fonseca, O. Salih, G.J. Kleywegt, A. Patwardhan, EMPIAR: the Electron Microscopy Public Image Archive, Nucleic Acids Res, 51 (2023) D1503–D1511.

